# *Bdellovibrio’s* Prey-Independent Growth is Fuelled by Amino Acids as a Carbon Source

**DOI:** 10.1101/2023.11.24.568592

**Authors:** C Herencias, MV Rivero-Buceta, S Salgado, F Baquero, R del Campo, J Nogales, MA Prieto

## Abstract

Identifying the nutritional requirements and growth conditions of microorganisms is crucial for determining their applicability in industry and understanding their role in clinical ecology. Predatory bacteria such as *Bdellovibrio bacteriovorus* have emerged as promising tools for combating infections by human bacterial pathogens due to their natural killing features. *Bdellovibrio’s* lifecycle occurs inside prey cells, using the cytoplasm as a source of nutrients and energy. However, this lifecycle supposes a challenge when determining the specific uptake of metabolites from the prey to complete the growth inside cells, a process that has not been completely elucidated. Here, following a model-based approach we illuminate the ability of *Bdellovibrio bacteriovorus* to replicate DNA, increase biomass, and generate adenosine triphosphate (ATP) in an amino acid-based rich media in the absence of prey, keeping intact its predatory capacity. In this culture, we determined the main carbon sources used and their preference, being glutamate, serine, aspartate, isoleucine, and threonine. This study offers new insights into the role of predatory bacteria in natural environments and establishes the basis for developing new *Bdellovibrio* applications using appropriate metabolic and physiological methodologies.

## INTRODUCTION

A complete understanding of the physiological and nutritional requirements that enable optimal cell growth plays an important role in the use of microorganisms in biotechnological, industrial and medical applications. In recent years, there has been an increase in new approaches to efficiently growing microorganisms, by taking advantage of new knowledge of their physiology [1, 2]. *Bdellovibrio bacteriovorus* is a gram-negative predatory bacterium belonging to the group of *Bdellovibrio* and like organisms (BALOs) that preys on other gram-negative bacteria. This species is ubiquitous, given that it can be found in soil, aquatic environments, and in human commensal microbiota [3]. Through a periplasmic predation strategy, *Bdellovibrio* uses the nutrients in its prey’s cytoplasm as carbon and energy sources for growth. Once the prey is exhausted, *Bdellovibrio* septates into its progeny and finally lyses the prey cell to begin another attack (**Fig. 1a**). Under certain conditions (limited nutrients/prey), certain *Bdellovibrio* strains can change their growth strategy and develop as prey-independent cells[4]. These *Bdellovibrio* variants can grow in the absence of prey bacteria, and their genotypic characterisation has revealed that they possess various mutations in genes related to prey recognition and attachment [5–10].

**Figure 1.**
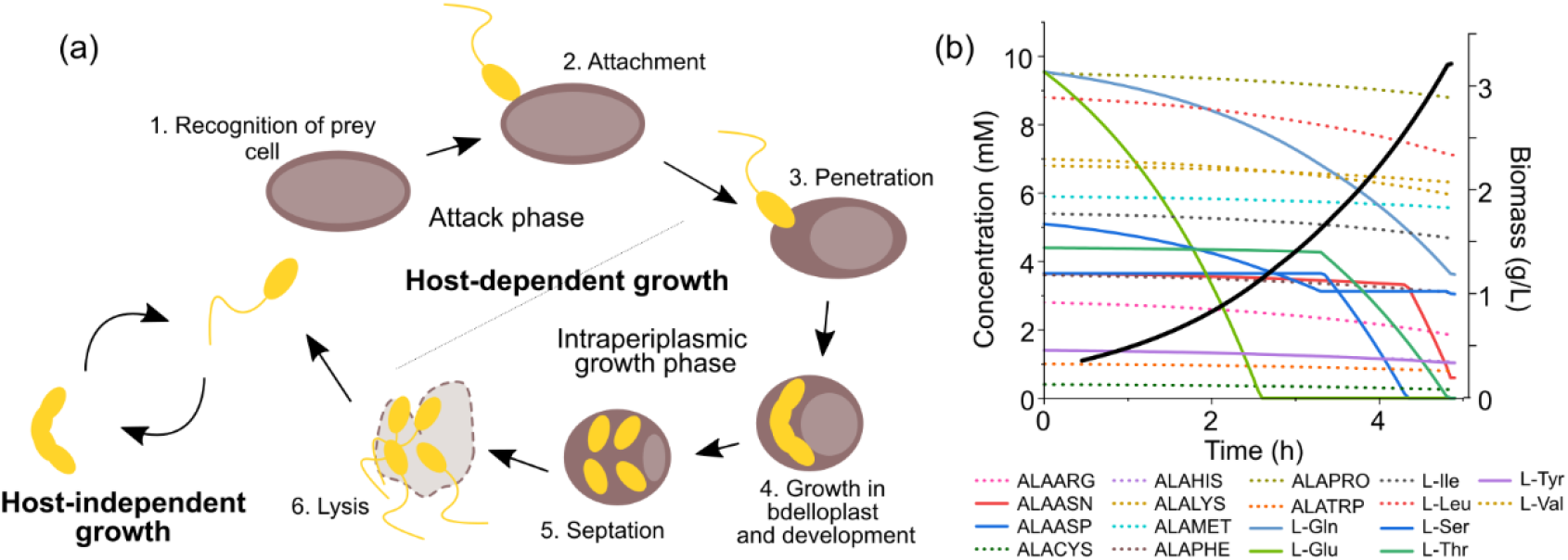
Representation of B. bacteriovorus lifecycle and metabolic simulation of its growth capacity. **a)** During host-dependent growth, B. bacteriovorus cells follow these steps: 1) Prey recognition: Bdellovibrio moves towards prey-rich regions. 2) Attachment: Bdellovibrio anchors to the host cell, which leads to the infection. 3) Penetration: Bdellovibrio enters the periplasm of the prey cell. 4) Growth in bdelloplast and development: the prey appears rounded due to cell wall modification, and Bdellovibrio grows in the periplasm and replicates its DNA. Bdellovibrio uses the prey biomolecules as a source of nutrients. 5) Septation: Bdellovibrio septates when resources become limited and matures into new individual attack phase cells. 6) Lysis: mature attack-phase cells lyse the cell wall of the bdelloplast, are released in the environment and start a new search for fresh prey. The complete host-dependent cycle takes approximately 4 h. During host-independent growth (left), Bdellovibrio cells elongate into filaments using the nutrients available in the environment to grow and finally sept into several daughter cells. **b)** Analysis of the growth capacity of B. bacteriovorus using the iCH457 model. Colour lines represent the prediction of amino acid consumption present in the rich media. The black line indicates the prediction of biomass production.

Derived from its lifestyle, multiple applications of the predators in various fields have been proposed, such as biocontrol agents (food preservation and water treatment), living antibiotics (to combat multiresistant pathogens), and hydrolytic enzyme producers[3, 11].

Several biochemical studies have highlighted aspects of the energetic metabolism of *Bdellovibrio* using diverse substrates such as amino acids and RNA[12–17]. Analysis of the *Bdellovibrio* genome revealed the presence of complete biosynthetic and degradation pathways for fatty acids; however, the lack of a phosphotransferase system makes the use of carbohydrates unlikely[18, 19]. More recent studies have examined the possibility of growing predator cells in a rich medium, offering new biotechnological perspectives[20]. Cell survival is minimally altered during prey starvation in the attack phase[20], and there is a slight increase in cell size and stimulation of protease secretion[21].

Currently, the precise nutritional requirements that allow for the growth of these predators are not entirely elucidated, mainly due to this organism’s host-dependent nature[11]. Analysing which metabolites from the prey cells are taken up during intraperiplasmic growth is a challenge, given the vast repertoire of molecules contained in the prey cell. We have reported a genome-scale metabolic model of this predatory bacterium (*i*CH457) to characterise and predict the metabolic potential and behaviour of the biphasic lifestyle of *B. bacteriovorus*[19]. Our model simulations have consistently postulated the growth capacity of *Bdellovibrio* in a rich amino acid medium in the absence of prey, based on its metabolic competencies.

The aim of the present study was to determine the growth and metabolic capabilities of the host-dependent *B. bacteriovorus* 109J, a laboratory reference strain, in a nutrient-rich but prey-free medium, and the uptake of metabolites during cellular growth. A better understanding of how these bacteria grow and their physiological and metabolic requirements will provide insight into their role in various microbial environments and increase the possibility of producing and applying predators rationally and safely in industry and clinical practice.

## MATERIALS AND METHODS

### In silico growth simulations

Biomass production and amino acids consumption were calculated using the dynamic Flux Balance Analysis (dFBA) of the COBRA toolbox 3.0 [22] in the environment MATLAB R2022a (MathWorks, Natick, MA, USA) using the Gurobi Solver (v9.5.2, Gurobi Inc.). Biomass was set as the objective function and the bounds of the exchange fluxes were set as the rich *in silico* medium as described previously [19]. Amino acids exchange reactions were set as “substrateRxns”, and “initConcetrations” of amino acid or dipeptides were based on Lysogeny Broth[23]. Since L-alanine is supplied in the form of dipeptides, the initial concentration of L-alanine was set in 0.001 mM, as well as the other unsupplied dipeptides, to avoid being set unlimited for the function dFBA.

### Strains, media and growth conditions

*B. bacteriovorus* 109J was routinely grown at 30ºC in Hepes buffer (25 mM Hepes amended with 2 mM CaCl_2_·2H_2_O and 3 mM MgCl_2_·3H_2_O, pH 7.8) or DNB liquid medium (consisting of 0.8 g L^−1^ NB (Difco™) (supplemented with 2 mM CaCl_2_·2H_2_O and 3 mM MgCl_2_·3H_2_O) with *Pseudomonas putida* KT2440 as prey[24, 25]. To remove the prey, the co-cultures were filtered twice through a 0.45 μm filter (Sartorius). The axenic growth of *Bdellovibrio* was carried out in PYE10 (10 g L^−1^ peptone and 10 g L^−1^ yeast extract) medium. CAV defined medium is composed of 200 μM of each amino acid: phenylalanine, glutamate, aspartate, threonine, serine, glycine, proline, isoleucine, leucine, valine and alanine. CAV medium was supplemented with a solution of trace elements (composition 1000 × 2.78 FeSO_4_·7H_2_O g L^−1^, 1.98 MnCl_2_·4H_2_O g L^−1^, 2.81 CoSO_4_·7H_2_O g L^−1^, 1.47 CaCl_2_·2H_2_O g L^−1^, 0.17 CuCl^2^·2H^2^O g L^−1^, 0.29 ZnSO_4_·7H_2_O g L^−1^), 5 μM of KH_2_PO_4_, 5 μM of K_2_HPO_4_ and 5 μM of NHCO_3_. All these compounds are suspended in Hepes buffer.

### *B. bacteriovorus* viability calculation

Predator viability was counted as plate forming units per milliliter (pfu mL^-1^). It was calculated from a culture performing serial dilution from 10^−1^ to 10^−7^ in Hepes buffer and developing on the lawn of prey after 48-72 h of incubation at 30°C by using the double layer method[24, 25]. Briefly, 0.1 mL of the appropriate dilution was mixed with an additional 0.5 mL of prey cell suspension of *P. putida* KT2440 pre-grown in NB and prepared in Hepes buffer OD_600_ 10, vortexed and plated on DNB solid medium.

### Biomass calculations

Cellular biomass, expressed in grams of the cell dry weight (g CDW) per liter, was determined gravimetrically as previously reported(38). Briefly, ten milliliters of culture medium were centrifuged for 15 min at 13000 x g at 4°C. Cell pellets and the supernatants were separated and subsequently freeze-dried for 24 h and finally weighed. The biomass calculation was obtained from the pellet weight.

### Measurements of ATP intracellular levels

Intracellular ATP levels were determined by an ATP bioluminescence assay kit (ATP Biomass Kit HS, BioThema) according to the manufacturer’s instructions. To measure the intracellular ATP, 1 mL of *B. bacteriovorus* cells was centrifuged for 15 min at 13000 x g and 4°C and the pellet was suspended in 1 mL of saline solution (0.85% NaCl). For each condition, four different experiments were carried out and two technical replicates were measured. To normalize the ATP intracellular values, the number of viable predator cells was also measured by the double-layer method.

### Quantification of *B. bacteriovorus* genome number

The abundance of the predator (number of genomes) was estimated by quantitative PCR (qPCR). A DNA fragment of 121 bp located in the coding region of the housekeeping gene bd2400(39) was amplified using the oligonucleotides Bd2400-1 (5′-GCGACTCCAGAACAGCAGATT) and Bd2400-2 (5′-GAATCCGCGGACTGCATTGTA) [26]. qPCR was performed using the SYBR Green (LightCycler® 480 SYBR Green I Master) technology in a LightCycler 480 Real-Time PCR system. Samples were directly analyzed from the culture (containing either prey and predator or predator alone), without DNA extraction. For the calibration curve, genomic DNA was purified from a filtered co-culture of *B. bacteriovorus* 109J using the IllustraTM bacteria genomicPrep Mini Spin Kit (GE Healthcare) following the instructions of the manufacturers. For the measurements, 200 μL of each sample was collected and stored at −20°C until analysis. Samples were initially denatured by heating at 95°C for 5 min, followed by 45 cycles of amplification (95°C, 10 s; test annealing temperature, 60°C, 10 s; elongation and signal acquisition, 72°C, 10 s). For quantification of the fluorescence values, a calibration curve was made using serial dilution from 5 to 5·10^−7^ ng of *B. bacteriovorus* 109J genomic DNA sample. qPCR was performed with triplicate samples from three independent biological experiments. The negative control was achieved with genomic DNA from *P. putida* KT2440 as a template. The results were analyzed using the 2^-ΔΔCt^ method and the genome number per mL can be calculated as follows [27, 28]: n° of genomes of Bd = (ng of DNA in the sample by qPCR)/(weight of the amplicon), being the weight of the amplicon of 1.24·10^−10^ ng. The calculations of concentration (genome mL^-1^) consider that the genome of the predator contains one copy of the bd2400 gene. Genes covering the whole genome were selected to validate the qPCR method and are listed in **Table S2 and Table S3**. To estimate the real predator cells using the qPCR method we correlated viable cell number and genome number. Then we calculate a linear equation: y = 0.0027·x + 2.1·10^8^ used to calculate the final data for genome number.

### Amplification of DNA and *hit* locus sequencing

PCR amplifications were performed in the buffer recommended by the manufacturer adding 0.05 μg of template DNA, 1 U of Phusion DNA polymerase and 0.4 μg of each deoxynucleotide triphosphate. Conditions for amplification were chosen according to the GC content of the oligonucleotides used, provided by the manufacturer (Sigma-Aldrich). DNA fragments were purified by standard procedures using the Gene Clean Turbo Kit (MP Biomedicals). PCR products were purified using the High Pure Purification Kit (Roche Applied Science). The oligonucleotides used to amplify and sequence the different hit-related genes are listed in **Table S4**.

### Analysis of extracellular metabolites by untargeted GC-TOF-MS

The supernatants of the cultures were collected, frozen at −80°C and lyophilized. All GC identifications were based on retention times and/or comparisons with commercially available standards. Moreover, the National Institute of Standards and Technology (NIST)[29] Mass Spectra Library based on a pre-defined matching criterium (similarity index ≥ 70%,[30]against mass spectral libraries was used.

Prior to analyses, samples were derivatized to increase the volatility of polar metabolites. For this purpose, 5-10 mg of the lyophilized supernatant, 300 μL of pyridine and 200 μL of BSTFA (with 1% TMCS) were added to each sample and heated with stirring at 70 °C for 1 h. Subsequently, the reaction mixture was transferred to the GC autosampler vials for further GC-TOF-MS analyses[31].

Agilent 7890A Gas Chromatography (Agilent Technologies) coupled to Waters Micromass GCT Premier Mass Spectrometer (coupled to COMBI-PAL-GC (EI)) (Waters Corporation, Milford) was used for separation and detection in the GC-TOF-MS setup. A GC column (ZB-5MSplus, Phenomenex) of 30.0 m x 250 μm x 0.25 μm was used. Helium was used as the carrier gas at a flow rate of 1.0 mL min^-1^. The split ratio for the injector was set to 1:10, with a total injection volume of 2 μL. Front inlet and ion source temperatures were both kept at 270°C. Oven temperature was set to equilibrate at 60°C for 1 min, before initiation of sample injection and GC run. After sample injection, the oven temperature was increased at a rate of 6°C min^-1^ to 325°C and held at 325°C for 3 min. The MS detection was operated in EI mode (70 eV) with a detector voltage of 1900 V. Full scan mode with a mass range of m/z 50–800 was used as the data acquisition method. The metabolite identities will be confirmed by targeted analysis.

### Phase-contrast microscopy

To monitor the growth of *Bdellovibrio* cells, cultures were routinely visualized using a 100X phase-contrast objective and images taken with a Leica DFC345 FX camera.

### Statistical analysis

Data sets were analyzed using Prism 6 software (GraphPad Software Inc.). Comparisons between two groups were made using Mann-Whitney-test followed by a no-parametric Krustal-Wallis analysis. The relation between the microbial counting and the genome number was analyzed by Kendall analysis. Data was represented using a R custom script and the ‘ggplot2’ package.

## RESULTS

### Rich medium supports the growth of *B. bacteriovorus* 109J in the absence of prey

The rational application of predatory bacteria for biotechnological or clinical purposes depends, to a large extent, on the level of understanding of the bacteria’s lifecycle and growth requirements. The biphasic lifecycle of *B. bacteriovorus* can complicate its cultivation at a large scale and its consequent applications; however, recent evidence of its metabolic capabilities for axenic growth will facilitate these processes[19]. In order to provide further insights on this topic, we employed the curated genome-scale model for *B. bacteriovorus* (*i*CH457) to predict cell growth by conducting a dynamic flux balance analysis (dFBA). Combining this approach with the optimization of biomass production under rich medium conditions [rich *in silico* medium[19]] it was predicted a specific growth rate of 0.486 h^−1^ (**Fig. 1b**). Furthermore, we found an interesting hierarchy in the amino acids consumption being glutamate the first carbon source used, followed by glutamine, serine, threonine, and asparagine (**Fig. 1b**).

The above *in silico* results strongly argue in favor of the full capacity of *Bdellovibrio* to grow axenically using amino acids as carbon and energy sources. To test this hypothesis, we cultivated the bacteria for 96 h in an amino acid-based media containing peptone and yeast extract (PYE10) and monitored growth parameters such as the viable cell count, the number of genomes, the intracellular ATP levels, and the biomass production. The results showed that while the number of viable predators decreased by 1 log_10_ after 96 h, the genome number increased by 0.5 log_10_, suggesting that a number of the surviving cells replicated their DNA without compromised viability (**Fig. 2a and b**). In other words, a portion of *B. bacteriovorus* cells can replicate DNA in PYE10 while others lose viability. The total biomass content and intracellular ATP concentration of the axenically grown *B. bacteriovorus* also increased up to 2.5-fold and 5-fold, respectively (**Fig. 2c and d**). Under these growth conditions, we obtained a homogenous population that would allow for a reliable and reproducible analysis of their physiological and metabolic states. Unexpectedly, the cells in the axenic culture were significantly larger than those in the control culture in the HEPES buffer (the mean size of PYE-Bd and HEPES-Bd were 1.684 and 0.6884 μm, respectively; t-test with Welch’s correction, p<0.001) (**Fig. 3**). In contrast, when the predators were inoculated in HEPES buffer, all monitored growth parameters decreased, indicating cell death.

**Figure 2.**
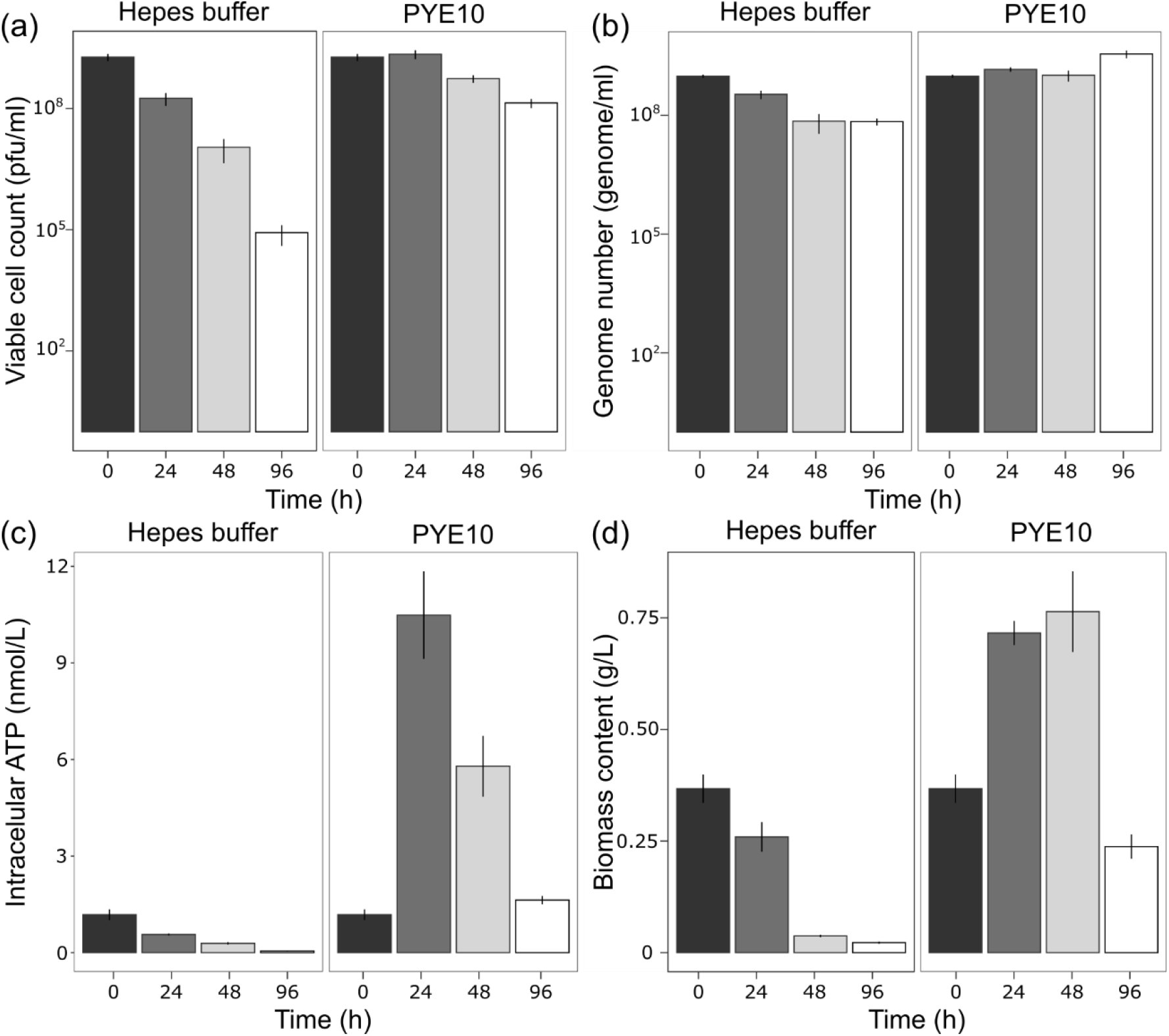
Growth parameters of B. bacteriovorus 109J growing in the absence of prey in HEPES buffer, PYE10 rich. **a)** Viable cell counts of Bdellovibrio measured as pfu/mL. **b)** Genome number measured using the bd2400 housekeeping gene. **c)** Measurement of intracellular ATP levels of Bdellovibrio cells. **d)** Total biomass content. Error bars indicate the standard deviation of three biological replicates. Statistical analyses are listed in **Table S1**.

**Figure 3.**
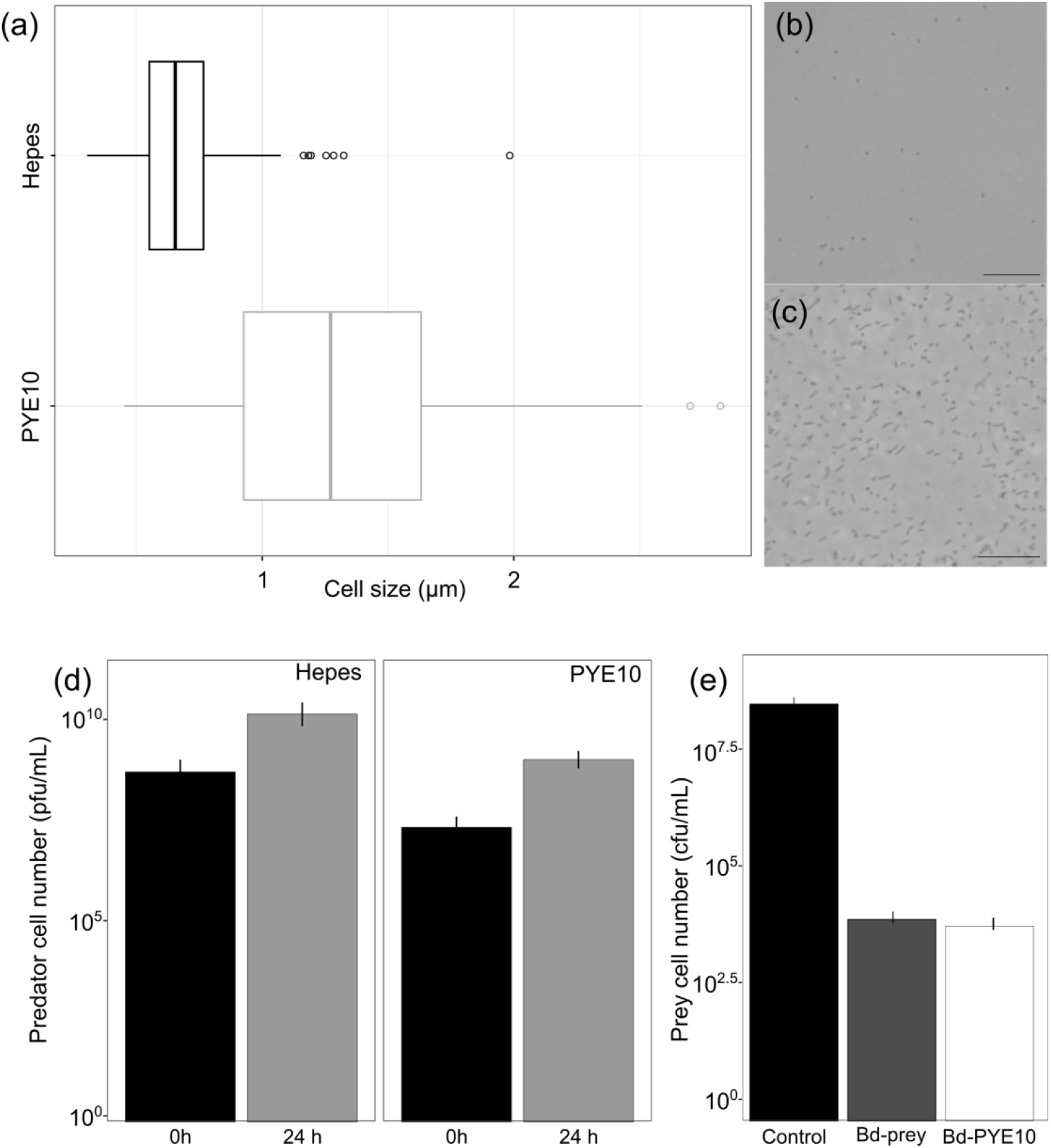
Morphology and predatory ability of B. bacteriovorus after growth in HEPES or PYE10 medium. **a)** Boxplot representations of the cell size population in μm using ImageJ software. Vertical lines within boxes indicate median values, left and right hinges correspond to the 25th and 75th percentiles, and whiskers extend to observations within 1.5 times the interquartile range (n=150, Mann-Whitney U test; p-value <0.01). **b) and c)** Phase-contrast micrography of Bdellovibrio growing in HEPES buffer and PYE10 medium, respectively. Black bar corresponded with 10 μm. **d)** Determination of predator cell counts after axenic growth in PYE10 medium or incubated in presence of prey cells. Error bars indicate the standard deviation of six biological replicates. **e)** Determination of prey viable cell count after predation of B. bacteriovorus 109J cells routinely grown with prey (Bd-prey) and prey viable number after predation with Bdellovibrio grown axenically (Bd-PYE10). Error bars indicate the standard deviation of six biological replicates.

To confirm the predatory capability of the *B. bacteriovorus* cells after incubation in prey-free PYE10 rich media, we performed predation experiments on *P. putida* KT2440. Our analysis of prey viability after 24 h of predation confirms the killing activity of the axenic *Bdellovibrio* cells, thereby ruling out the possibility of loss of the predatory genotype after cultivation in prey-free conditions (**Fig. 3**).

The genotype of the wild-type *B. bacteriovorus* 109J was also verified by sequencing the genes responsible for the host-independent (HI) phenotype (bd0108, bd3461, and bd3464 [32]). Sequences were compared with two HI *Bdellovibrio* mutants, HI18 and HI24 (Prof. Jurkevitch lab collection), and no significant mutations were accumulated during the growth in the absence of prey (**Fig. S1**). As a whole, these results demonstrated the active metabolism of the wild-type predator grown in PYE10 media, supported by the active DNA replication, the increased biomass content and intracellular ATP concentration, the increased cell size, and maintenance of the predatory capability.

### Analysis of metabolite consumption during *Bdellovibrio* culture in PYE10 medium

To determine the nutrient composition of the PYE10 medium and monitor the uptake of metabolites by *B. bacteriovorus*, we conducted a gas-chromatography/quadrupole time-of-flight (GC-QTOF) analysis. Predator cells were incubated in PYE10 medium; after 96 h, the supernatant was collected and analysed. Amino acids constituted 75% of all nutrients present and detected in the PYE10 medium (**Fig. 4a**). In addition, the main non-amino acid metabolites detected in the PYE10 medium were compounds related to secondary metabolism such as butanoic, propanedioic, and gluconic acids. The analysis also revealed relatively high amounts of trehalose, xanthine, and pyranose derivatives and phthalic acid (**Fig. S2 and Supplementary Dataset)**. There were minimal spontaneous variations in the composition of the control medium without predator throughout the experiment, as previously reported[33]. In contrast, the presence of *Bdellovibrio* drastically altered the composition of the medium, primarily through the consumption of amino acids (**Fig. 4b, Fig. S3, and Supplementary Dataset**). The relative amounts of glutamate, serine, aspartate, isoleucine, and threonine were significantly lower compared with the control culture at the start of the experiment (unpaired t-test <0.02). The results, highly agreeing with *in silico* predictions confirmed the preference consumption of amino acids over small organic acids also present in the medium.

**Figure 4.**
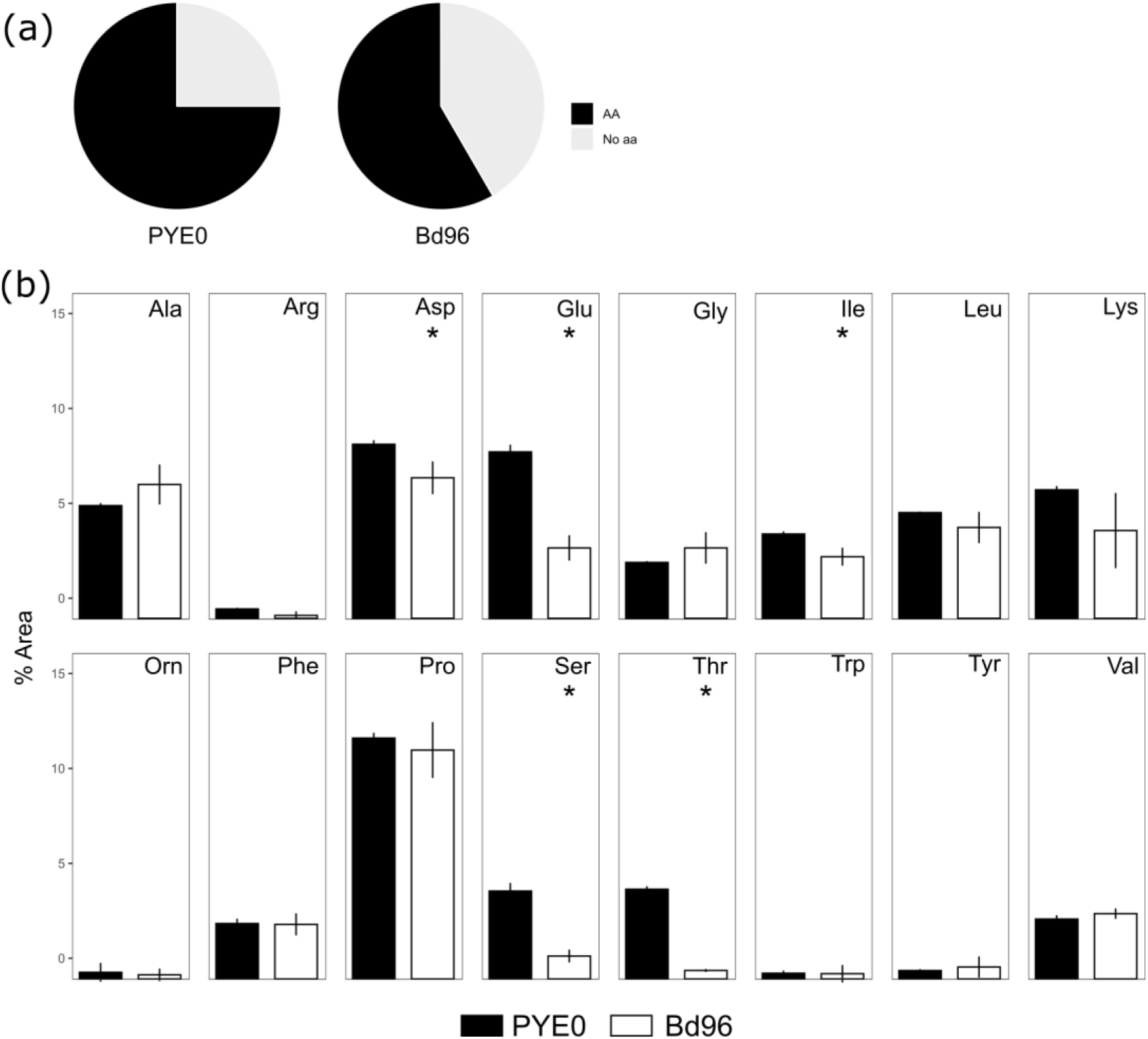
Relative composition of PYE10 medium during Bdellovibrio axenic growth. **a)** Percentage of amino acid and non-amino acid compounds of PYE10 medium at the beginning of the experiment and after 96 h of Bdellovibrio growth. **b)** Amino acid consumption during Bdellovibrio axenic growth (Bd 96 h) compared with control (PYE0). Error bars indicate the standard deviation of three biological replicates. Ala, alanine; Arg, arginine; Asp, aspartic acid; Glu, glutamate; Gly, glycine; Ile, isoleucine; Leu, leucine; Lys, lysine; Orn, ornithine; Phe, phenylalanine; Pro, proline; Ser, serine; Thr, threonine; Trp, tryptophan; Tyr, tyrosine; Val, valine. Asterisks indicate statistically significant differences compared with time PYE0 (Unpaired t-test <0.02).

### Growth of *B. bacteriovorus* using amino acids as a carbon and nitrogen source

A key limitation of the growth rate of bacteria is nutrient concentration. The above results suggest the use of amino acids is sufficient to support *B. bacteriovorus* growth. To validate this hypothesis, we designed a medium, which is composed only of amino acids CAV medium (see Materials and Methods). To monitor the growth dynamics of *B. bacteriovorus* in the absence of prey, we incubated the predator cells in the defined CAV medium. The growth was monitored by measuring the viable cell count, the genome number, the biomass content, and the intracellular ATP concentration. The analysis of these parameters (**Fig. 5**) revealed maintenance of an active metabolism. There was no increase in the viable cell count or genome number under this condition, but the biomass content increased significantly during the 48 h (p<0.03, Krustal-Wallis t-test). After this point, cells strongly sense starvation, resulting in cell energy depletion and biomass reduction at 96 h.

**Figure 5.**
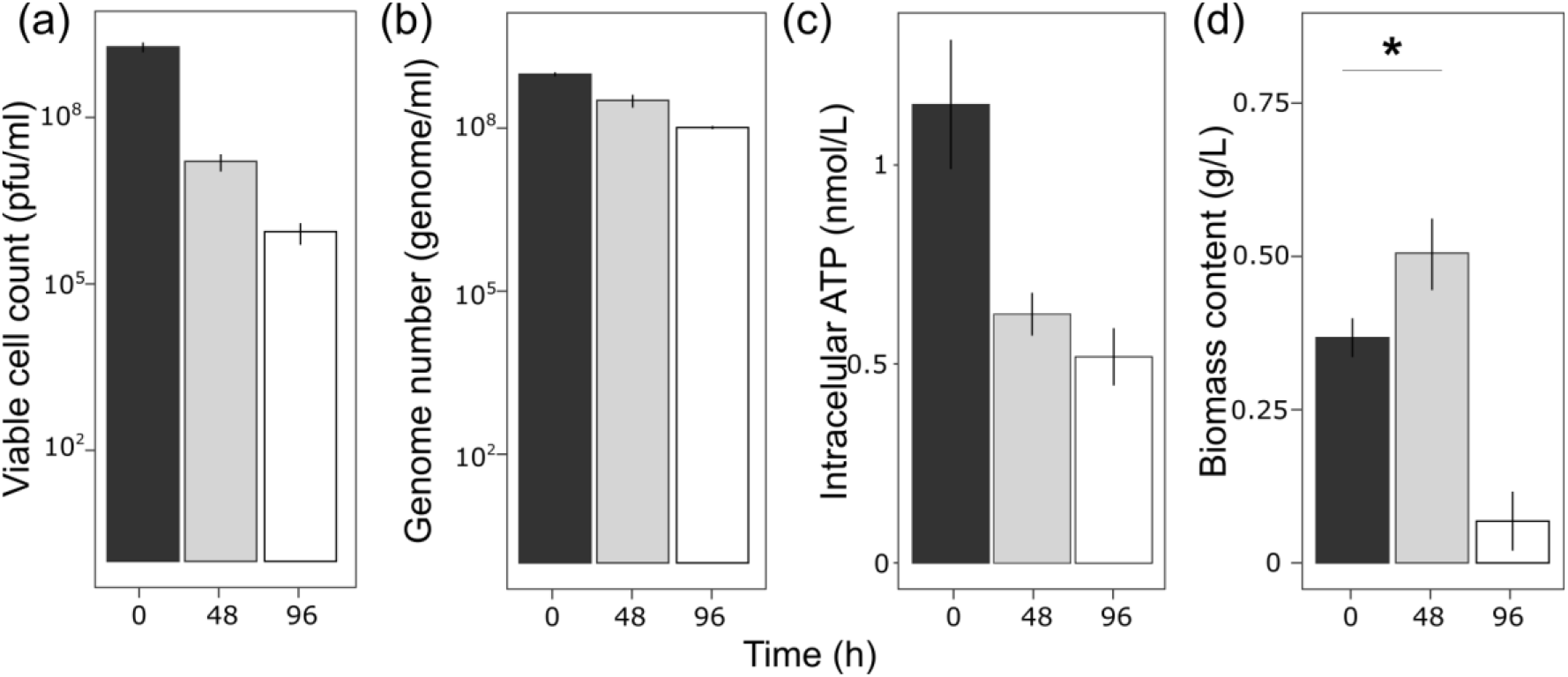
Growth parameters of B. bacteriovorus 109J growing in CAV medium. **a)** Viable cell number of Bdellovibrio measured as pfu/mL. **b)** Genome number measured using the bd2400 housekeeping gene. **c)** Measurement of intracellular ATP levels of Bdellovibrio cells. **d)** Total biomass content. Error bars indicate the standard deviation of three biological replicates. Asterisks indicate statistically significant differences compared to time 0 (Krustal-Wallis t test <0.03).

## DISCUSSION

The evolution of biological systems is characterized by their ability to acquire diverse strategies to adapt to various environments. A significant aspect of this adaptation is the preference for specific carbon and energy sources, which is a result of metabolic adjustments to the environment. *B. bacteriovorus* has evolved to thrive in the periplasmic space of the prey cell by using the prey cell’s cytoplasmic components[11, 34]. Several reports have contributed valuable insights into the energetic metabolism and growth conditions of *Bdellovibrio*[35, 36]; however, there is still a dearth of evidence regarding the specific nutritional requirements of *Bdellovibrio* and preferred carbon source as a predator. This study analysed the carbon and energy requirements necessary to support the growth of *Bdellovibrio*. Understanding these critical factors is vital for potential industrial and medical applications of *Bdellovibrio*.

The bacterial cytoplasm serves as a highly concentrated compartment, containing a significant portion of the cell’s macromolecules (30–40%) and proteins (over 70%)[37]. This abundance makes the cytoplasm an exceptionally nutrient-rich environment that facilitates the growth of *B. bacteriovorus*[38, 39]. Most of the cytoplasmic proteins are crowded into the vicinity of the cellular envelope, and prey cellular deformation after the bdelloplast formation likely concentrates and increases accessibility to the protein layer[40]. *Bdellovibrio* proteases can permeate the altered envelope and use these proteins as substrates to obtain small peptides and amino acids. Based on its genome analysis, *Bdellovibrio* exhibits a wide array of systems dedicated to the uptake of amino acids in the form of di- or tri-peptides[41].

Our research validated the functional metabolism of *B. bacteriovorus* under axenic growth conditions predicted by metabolic modelling, specifically using the nutrient-rich medium PYE10, in which the primary ingredients are amino acids. The results of our study confirmed the activation of DNA replication (genome number), ATP generation, and biomass production by *Bdellovibrio* under these conditions (**Fig. 2**). Additionally, the study substantiates the uptake of amino acids from PYE10 through GC-MS analysis (**Fig. 4**).

The predator, *Bdellovibrio*, is commonly described as host-dependent, which poses a challenge to its efficient application and control. Our findings reveal that *Bdellovibrio* can be cultured as host-independent bacteria while maintaining its effective killing efficiency (**Fig. 3**). Even after prolonged starvation conditions without prey, predator cells can reduce the prey population as efficiently as in the *Bdellovibrio* control culture during the attack phase. This remarkable result highlights the role of *Bdellovibrio* as a regulator of bacterial populations, not only in aquatic and soil environments [likely invading protein-rich bacterial aggregates or biofilms attached to microbiotic particles[42]] but also in the human commensal microbiota[43, 44]. The predator’s survival therefore does not solely rely on individual predation events.

Furthermore, our experiments demonstrated that incubation in the defined minimal medium CAV sustained an active metabolism in *Bdellovibrio* (**Fig. 5**). Amino acids are precursors of biomass components that fuel the bacterial metabolic network at various points[45]. For instance, glutamate that enters from α-ketoglutarate plays a key role in anaplerotic reactions. Notably, glutamate dehydrogenase directly fuels the tricarboxylic acid cycle by producing α-ketoglutarate, a reaction that serves as an excellent source for reducing equivalents, which are essential metabolites for anabolic reactions. Serine, which enters from 3-phosphoglycerate, supplies the upper path of glycolysis. Leucine and isoleucine enter from pyruvate, connecting with the fatty acid synthesis. This study experimentally validated and confirmed the functional capabilities of the *Bdellovibrio* proteolytic machinery and the key role of amino acids in sustaining efficient growth, with or without prey.

In summary, the axenic cultivation of *B. bacteriovorus* under controlled conditions and the modulation of medium composition (PYE10 or CAV) ensures efficient predator metabolism and growth. Although extensive studies are needed before predator biotechnological and clinical applications become available, the findings presented here open new avenues of research aimed at understanding the nutritional requirements of *B. bacteriovorus*.

## ACKNOWLEDGMENTS

We appreciate the technical support of Ana Valencia. We are indebted to Prof. Edouard Jurkevitch (The Hebrew University of Jerusalem, Israel) for sharing the host-independent *Bdellovibrio* HI18 and HI24 used in this study.

