## Supplementary material for "*Bdellovibrio’s* Prey-Independent Growth is Fuelled by Amino Acids as a Carbon Source": Supplmentary Material

[1]Department of Microbiology, Hospital Universitario Ramón y Cajal, Instituto Ramón y Cajal de Investigacion Sanitaria (IRYCIS), Madrid, Spain. [2]Polymer Biotechnology laboratory, Biological Research Center-Margarita Salas, CSIC, Madrid, Spain. [3]Systems Biotechnology Group, Department of Systems Biology, Centro Nacional de Biotecnología, CSIC, Madrid, Spain, [4]Centro de Investigación Biomédica en Red de Epidemiología y Salud Pública, Instituto Carlos III, Madrid, Spain, [5]Centro de Investigación Biomédica en Red de Enfermedades Infecciosas-CIBERINFEC, Instituto de Salud Carlos III, Madrid, Spain. [6]nterdisciplinary Platform for Sustainable Plastics towards a Circular Economy-CSIC (SusPlast-CSIC), Madrid, Spain

[†]These authors contributed equally and should be considered as both first authors

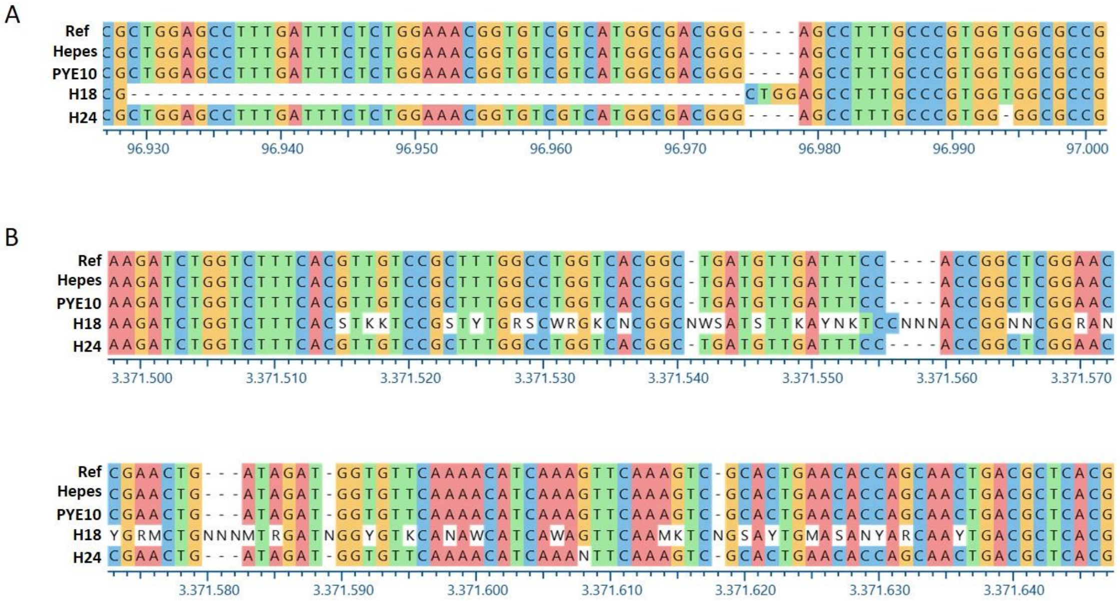


***Supplementary Figure 1. DNA hit loci sequences of* B. bacteriovorus *growing PYE10 medium. a)*** *bd0108 gene and* ***b)*** *bd3461 gene. These sequences were aligned with CLUSTALX (Thompson et al., 1997) and processed in Geneious (Geneious 10.2.2 (https://www.geneious.com)) to identify conserved regions. Sequences are presented in the 5'→3' orientation. Ref, reference sequence of* B. bacteriovorus *109J available in National Center for Biotechnology Information database; HEPES, sequence of* Bdellovibrio *control cells; PYE10, sequence of* Bdellovibrio *cells growing in PYE10 medium; HI18, sequence of HI18 prey-independent strain; HI24, sequence of HI24 prey-independent strain. Colours indicate different nucleotides.*

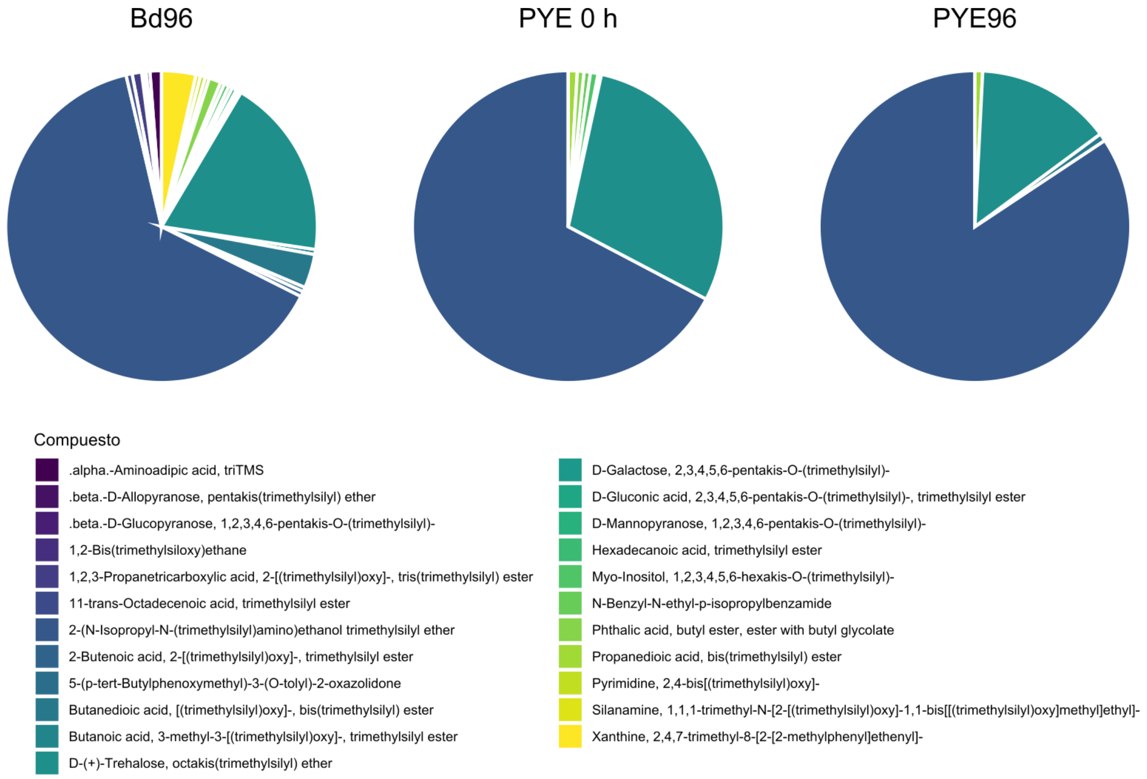


***Supplementary Figure 2. Relative abundance of metabolites detected by GC-QTOF.*** *The pie chart shows 23 metabolites related to organic acids and derivatives.*

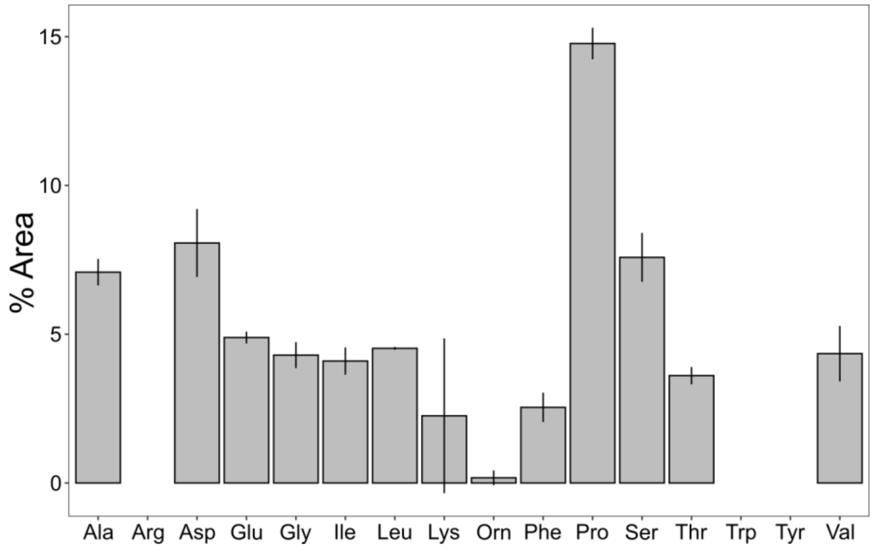


***Supplementary Figure 3. Change in amino acid composition of PYE10 medium after 96 hours.*** *Error bars indicate the standard deviation of three replicates. Ala, alanine; Arg, arginine; Asp, aspartic acid; Glu, glutamate; Gly, glycine; Ile, isoleucine; Leu, leucine; Lys, lysine; Orn, ornithine; Phe, phenylalanine; Pro, proline; Ser, serine; Thr, threonine; Trp, tryptophan; Tyr, tyrosine; Val, valine.*

**Supplementary Table 1: p-values of Mann-Whitney test of *Bdellovibrio´*s growth parameters**

| | Viable count | Genome count | ATP measurement | Biomass content |
|---|---|---|---|---|
| **Hepes 0h-Hepes 96 h** | 0.002 | 0.002 | 0.002 | 0.01 |
| **PYE10 0h-PYE10 96h** | 0.002 | 0.004 | 0.04 | 0.03 |
| **Hepes 96h-PYE 96** | 0.002 | 0.002 | 0.002 | 0.03 |
| **CAV 0h-CAV 96h** | 0.01 | 0.01 | 0.01 | 0.01 |
| **Hepes 96h-CAV 96h** | 0.03 | ns | 0.01 | ns |
| **PYE 96h- CAV 96h** | 0.01 | 0.01 | 0.01 | ns |

**Supplementary Table 2. Genes and oligonucleotides used to validate the qPCR method to calculate the genome number.** Locus on the genome: *bd2478* (293927-292831), *bd0562* (170745-171884), *bd1803* (339602- 337938) and *bd3230* (22311-23105)

| Gene | Protein | Gene Length (pb) | Amplicon Mw (g mol $^{-1}$) | Oligonucleotide 5´-> 3´ |
|---|---|---|---|---|
| ***Bd2478*** | Alanine dehydrogenase | 1107 | $1.51 \cdot 10^5$ | F: CAGCTTCGCTTCTGCAGCCAAGTG<br>R: GCGTGCGTCAGTATGTGAGCGAAG |
| ***Bd0562*** | Citrate synthase | 1140 | $1.04 \cdot 10^5$ | F: GCTGGCGGATTCTCCACTCAAC<br>R: GCTTCAGCGCCGTTGTCGTTGG |
| ***Bd1803*** | Fatty acid CoA ligase | 1665 | $8.22 \cdot 10^4$ | F: GGTTGCAGCACACAACTGGAGAAGC<br>R: CGTGTGAAAGTCAGCGTGGCAGG |
| ***Bd3230*** | Thymidilate synthase | 785 | $1.17 \cdot 10^5$ | F: GGCCGCACCATCGATCAGATCAC<br>R: CCAGCTGTATCAGCGCAGTGCTG |

**Supplementary Table 3. Genome number calculation of *B. bacteriovorus* using different reference genes.**

| Gene | Sample | Genome number (genome $mL^{-1}$) | Standard deviation | Predicted viable (pfu $mL^{-1}$) |
|---|---|---|---|---|
| ***Bd2400*** | 0 h | 6,36E+09 | 7,31E+08 | 1,72E+07 |
| | 24 h | 1,06E+12 | 1,04E+10 | 2,87E+09 |
| ***Bd0562*** | 0 h | 5,55E+09 | 8,14E+08 | 1,50E+07 |
| | 24 h | 1,23E+12 | 1,34E+10 | 3,33E+09 |
| ***Bd2478*** | 0 h | 7,59E+09 | 1,49E+09 | 2,05E+07 |
| | 24 h | 1,29E+12 | 1,11E+10 | 3,49E+09 |
| ***Bd1803*** | 0 h | 3,82E+09 | 7,72E+08 | 1,03E+07 |
| | 24 h | 9,43E+11 | 6,63E+09 | 2,55E+09 |
| ***Bd3230*** | 0 h | 4,49E+09 | 8,08E+08 | 1,21E+07 |
| | 24 h | 7,97E+11 | 1,03E+10 | 2,15E+09 |

**Supplementary Table 4. Oligonucleotides sequences used to amplify the *hit* locus genes to corroborate the wild type genotype of the predator cells growing in rich medium.**

| Target gene | Oligonucleotide | Sequence 5´-> 3´ |
|---|---|---|
| ***Bd0108*** | Bd9 | CTAGCTAGCAGAAGGTGATTATATGAAAA |
| | Bd32 | AGGAAGCTTTACTGTCTTCCAGTCCCGGCTTTC |
| | BdRa | AGTGTATTGCCTACCGTACCG |
| | BdFa | GCCTCATTAGGGTCTTCGCC |
| ***Bd3461*** | 3461Ra | GACAAGAGGGTGGTCCGAAG |
| | 3461Fa | CTCTGTCACCACCACGTCC |
| | 3461Fb | CCAGCACGTTCATGTCGTTTTC |
| | 3461Fc | TTGTCGTAACCTGTACCGCC |
| ***Bd2400*** | Bd2400Ra | CGACTTGCAAACATGGACGT |
| | Bd2400Fe | TTCAACTGTTCTGGCGTTGC |
| | Bd2400Fd | CGAGACACTTCCACCCAGTT |
| | Bd2400Fc | GTTTCACTTGGTCCGCACAG |
| | Bd2400Fb | TTCACCATCCAGTCGTAGCG |
| | Bd2400fa | ATCAAACCCACACCCAGGAC |

**Supplementary Dataset.** This file includes all the raw data form the QTOF- MS analysis used in this study.
